# Expression-Based Inference of Human Microbiome Metabolic Flux Patterns in Health and Disease

**DOI:** 10.1101/2020.01.09.900761

**Authors:** Yiping Wang, Zhenglong Gu

## Abstract

Metagenomic sequencing has revealed that the composition of the gut microbiome is linked to several major metabolic diseases, including obesity, type 2 diabetes (T2D), and inflammatory bowel disease (IBD). However, the exact mechanistic link between the gut microbiome and human host phenotypes is unclear. Here we used constraint-based modeling of the gut microbiome, using a gene-expression based algorithm called FALCON, to simulate metabolic flux differences in the microbiome of controls vs. metabolic disease patients. We discovered that several major pathways, previously shown to be important in human host metabolism, have significantly different flux between the two groups. We also modeled metabolic cooperation and competition between pairs of species in the microbiome, and use this to determine the compositional stability of the microbiome. We find that that the microbiome is generally unstable across controls as well as metabolic microbiomes, and metabolic disease microbiomes even more unstable than controls.

## 2 Introduction

Metabolic flux is one of the most important phenotypes that can be measured for a cell [1]. It determines what types of nutrients the cell will uptake, and how it will use them to harvest energy and create the building blocks of a cell such as nucleic and amino acids [2]. This determines the growth rate of a cell and ultimately its fitness in an environment with a limited supply of nutrients. A large part of the cell’s regulatory machinery is therefore devoted to regulating metabolism [3], and changes in metabolism play an important role in the development of many diseases, such as cancer [4], diabetes [5], and heart disease [6].

In recent years, another area where metabolism has been shown to be important is the human gut microbiome, which has seen great interest for its impact on nearly every aspect of health of its host [7]. It has been shown that the gut contains a variety of microorganisms, almost all bacterial, which in total have roughly 10 times the cell count of the human host, and 100 times as many genes [8]. The gut microbiome carries out multiple important functions for its human host, including digestion of fibers that can contribute up to 10% of the body’s energy needs, synthesis of essential vitamins, and competition with pathogens that could infect humans [9]. Not surprisingly, then, disruptions in the microbiome’s composition have been linked to numerous diseases, including gut diseases such as colon cancer and Crohn’s disease [10], as well as many that may seem unrelated to it, such as atherosclerosis [11]. Typically, these links are inferred by a technique similar to GWAS, by measuring species composition based on sequencing of the 16s rRNA gene, and comparing this composition between healthy and diseased individuals [12].

However, even though microbiome composition may be clearly associated with disease, it is still unclear whether there is any association with changes in microbiome metabolic flux. Given the crucial role of gut-derived metabolites, such as SCFAs [13], in the human body, it seems reasonable to assume that changes in gut microbiome metabolic flux play an important role in the development of certain diseases. One of the most striking findings from the recently completed Human Microbiome Project is that healthy subjects may have very different gut microbiome compositions, but functionally had almost the same metagenomic coverage of metabolic pathways [14]. However, metabolic flux also remains one of the most difficult phenotypes to experimentally measure. The current state-of-the-art is to use 13C isotopes to label metabolites in the cell, followed by mass spectrometry to measure the concentration of labeled metabolites. The overall labeling patterns can be used to computationally infer flux [15]. These methods suffer from two major limitations. First, mass spectrometry cannot accurately measure low-abundance labeled metabolites, which may be crucial for accurate flux determination on a genomic scale [16]. Second and more importantly, mass spectrometry measurements usually involve an average across a sample of cells [16], involving many different species in the gut microbiome, and so cannot resolve the metabolic state of individual species.

To overcome these challenges, several previous groups [17] [18] have used a class of algorithms called constraint-based modeling (CBM) [19] to computationally infer metabolic flux in the microbiome. As described further in the Methods section, CBM methods are designed to take a minimal set of physical and chemical assumptions (constraints) about metabolic flux, and use these to determine a reasonable solution space of metabolic flux distributions. However, in order to deal with the presence of multiple species in the gut microbiome, these previous methods must make additional assumptions about metabolic interactions between species, also described further in Methods. We have therefore developed two innovations in our paper to more accurately model gut microbiome metabolism. First, we applied a previously developed CBM method in our lab called FALCON [20]. FALCON takes the solution space determined as above, and then determines one particular, optimal flux distribution, by optimizing the agreement between the metabolic flux of each reaction, and the enzymatic expression associated with that reaction. This approach follows the general principle that highly active reactions require large amounts of expressed enzyme in order to function, and vice versa [21]. We believe that FALCON requires fewer assumptions than previous work, and therefore may be more suited to modeling microbiome metabolism.

Second, we also used three different types of metabolic network models for the microbiome, each of which captures different types of important interactions. As described below, we first used the single species AGORA models [22], a set of 773 metabolic reconstructions for individual gut microbiome species. The strength of these models is that they allow us to analyze major flux differences between samples, and break down which individual species are responsible for these changes. However, a drawback of the single species models is that they cannot capture any metabolic interactions between species. To capture these, we also developed a merged gut microbiome model, in which all reactions in the AGORA models are connected into a single network of 3670 reactions. Although the merged model loses all species specificity, it allows modeling of metabolic interactions among any number of microbiome species, as all reactions are included within it. Finally, between single-species and merged models, we also examined all pairwise models, in which all possible pairs of AGORA models in a sample are placed together in a common environment. This allows us to capture a subset of metabolic interactions, those between all pairs, while still retaining species specificity. As will be seen, the pairwise models are also sufficient to investigate the compositional stability of the gut microbiome.

We applied FALCON to analyze the metagenomic expression data for microbiomes from patients of three different metabolic diseases, namely obesity [23], Type 2 diabetes [24], and IBD [25], using each of the three models we have described. We calculated the resulting flux distributions in all cases, and used the most appropriate model to determine the pathways which carry different amounts of flux between normal and obese subjects, as well as stability differences between them. A flow chart of our workflow is shown in Figure 1.

**Figure 1:**
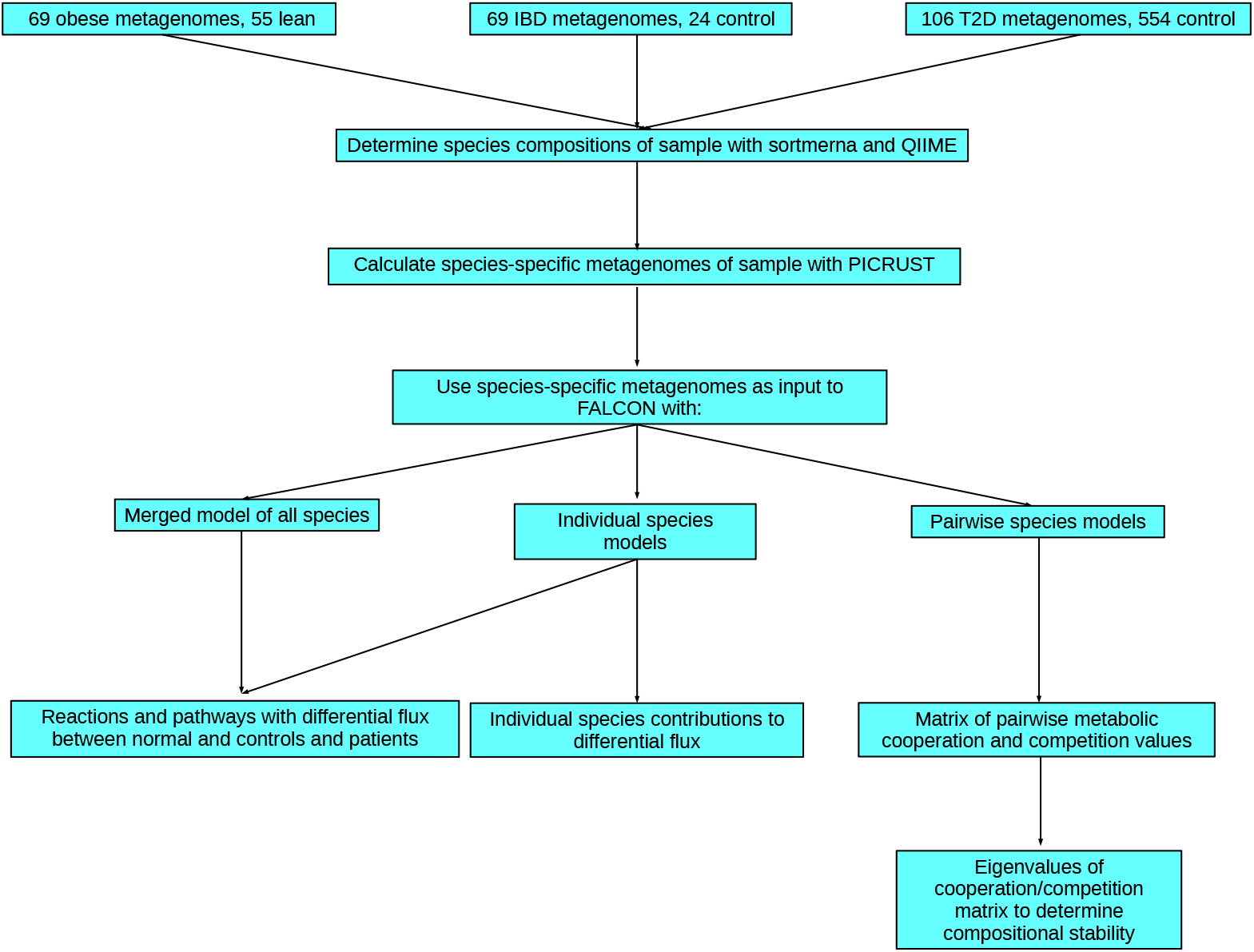
Flowchart of analyses carried out in this paper.

## 3 Methods

Metabolic networks can be considered as bipartite networks, in which each reaction is linked to the metabolites that it consumes and/or produces [19]. CBM methods thus model a metabolic network as a stoichiometric matrix *S*, in which each row represents a metabolite, each column a reaction, and at each row-column intersection is a coefficient representing how many molecules of a metabolite are involved in each reaction. Using this matrix, two major physico-chemical constraints may be imposed. First, at steady-state, the concentrations of metabolites in a cell are neither rising or falling. Therefore, given a vector of fluxes defined as *v*, this constraint may be written as the equation *S* · *v* = 0. Second, all metabolic reactions have a maximum rate, given that a cell can produce only a finite amount of enzymes, and enzymes’ efficiency is limited. This constraint may be written as *v* <= *v_max_*, where *v_max_* is the maximum rate of a metabolic reaction.

In the case of some reactions, *v_max_* can be inferred based on maximum levels of experimentally measured flux rates [2]. This is especially important for reactions which model the uptake of nutrients such as oxygen or glucose, which are often the limiting factor that constrains cell growth rates. For other reactions, *v_max_* is set to an arbitrary large value, usually +− 1000. This is done so that flux through these reactions will not become the limiting factor in cell growth rates [19].

Traditionally, the distribution of in a metabolic network inferred using flux balance analysis with a biomass objective. A biomass objective is a special reaction in the network represents the proper ratios of amino acids, nucleotides, and other metabolites necessary to create biomass. A flux distribution that maximizes the rate of the biomass reaction is considered to maximize cell growth rate, and therefore fitness. However, biomass optimization may not be the most accurate way to model metabolism in the microbiome, for several reasons. It is known that the human host can alter the composition of the microbiome through mechanisms such as immune targeting, and provision of nutrients like mucins to certain species [26]. Some bacterial species may follow strategies besides biomass optimization to survive in the microbiome, such as producing toxins that reduce growth of competitors [26]. Finally, in any simulation method involving biomass optimization in the microbiome, different species would be expected to have different biomass growth rates. This implies that species which grow the fastest would be able to outcompete other species in the microbiome, and eventually dominate the gut environment. As this is not observed experimentally, previous methods that simulate gut metabolism with FBA must incorporate additional constraints to prevent unbalanced growth rates [18] [17].

To avoid these problems, we instead chose to apply FALCON, which is a previously developed method in our lab that infers flux based on gene expression, by optimizing the correlation between flux and gene expression. Details of the FALCON algorithm may be found in [20]. Biomass rate is not optimized under FALCON, and indeed, in our FALCON simulations, we have never observed flux through the biomass reaction. However, the rationale behind FALCON is that high enzyme expression generally correlates with high flux, and vice versa [21]. Although this correlation is very weak in some cases, and has not been experimentally measured in the gut microbiome, we still believe that FALCON may provide a more principled simulation method than traditional FBA.

We furthermore applied FALCON to three different types of metabolic models for the gut microbiome, all of them based on the previously published AGORA database of gut microbiome models. The AGORA models are a set of 773 metabolic reconstructions for gut microbiome species. Importantly, they contain a set of estimated lower and upper bounds on uptake of common nutrients, such as glucose and amino acids, that are representative of a Western diet [22]. We used these limits for all simulations in our study.

Each of the three types of AGORA-derived metabolic models has their own strengths and drawbacks. First, we applied FALCON to the individual species-specific models from AGORA. This setting represents how metabolic flux would be distributed in each species if they were growing alone. Although simulations with these models do not capture any species interactions, they do show whether any particular species has significant metabolic differences between normal control and metabolic disease samples. They may also be useful to study the metabolism of individual microbiome species which can be cultured. Second, we applied FALCON to pairwise models, in which we modeled every possible pair of species, placed together in a common environment, i. e. sharing common nutrient uptake and waste product secretion, and then simulated with FALCON. As will be seen later, this is the most efficient way for us to elucidate all pairwise interactions among species, which can be used to calculate the compositional stability of the microbiome as a whole.

Finally, we merged all microbiome models in AGORA together, by combining all unique metabolic reactions across all species into one model, to form a very large merged model of 3670 reactions, representing the combined metabolic potential of the gut microbiome. The advantage of using this model is that it is able to capture how metabolic fluxes in the entire microbiome are distributed, taking into account all possible species interactions. That is, individual species in the gut may strongly interact with each other in groups of three or more species, and it is impossible to individually model all such possible groups. However, the merged model allows all reactions in all species to be linked to each other, allowing interactions across the entire microbiome to be modeled.

Furthermore, the size of the merged model depends on which species are included, and it should be noted that the size quickly saturates as more individual models are included. This is shown in the left half of Figure 2, which shows the number of reactions in the merged model across 1000 randomized trials, in which models are selected from the AGORA set and added in a random order. The result is important, as the AGORA set does not cover all known gut microbial species, and potentially new species added to the model may add new reactions. We do not expect this to be a major drawback though, as such new reactions would only comprise a small fraction of the merged model’s total. Furthermore, the metagenomic samples we consider may have very different species compositions, and thus the number of metabolic models that can be imputed to each model may be very different, as shown in the right half of Figure 2. This figure shows the number of individual metabolic models present within each sample of our diabetes dataset. However, similar to the findings of the Human Microbiome Project [8], we would also expect that each sample should have very roughly constant coverage of metabolic pathways, as all microbiomes must carry out the same set of metabolic functions. Indeed, when we constructed the merged models for each of our diabetes samples, by merging all individual models in the sample, we found that they all contained almost the same number of reactions. We interpret this to mean that all samples do have complete coverage of all metabolic reactions in the microbiome, as expected from the result in Figure 2

**Figure 2:**
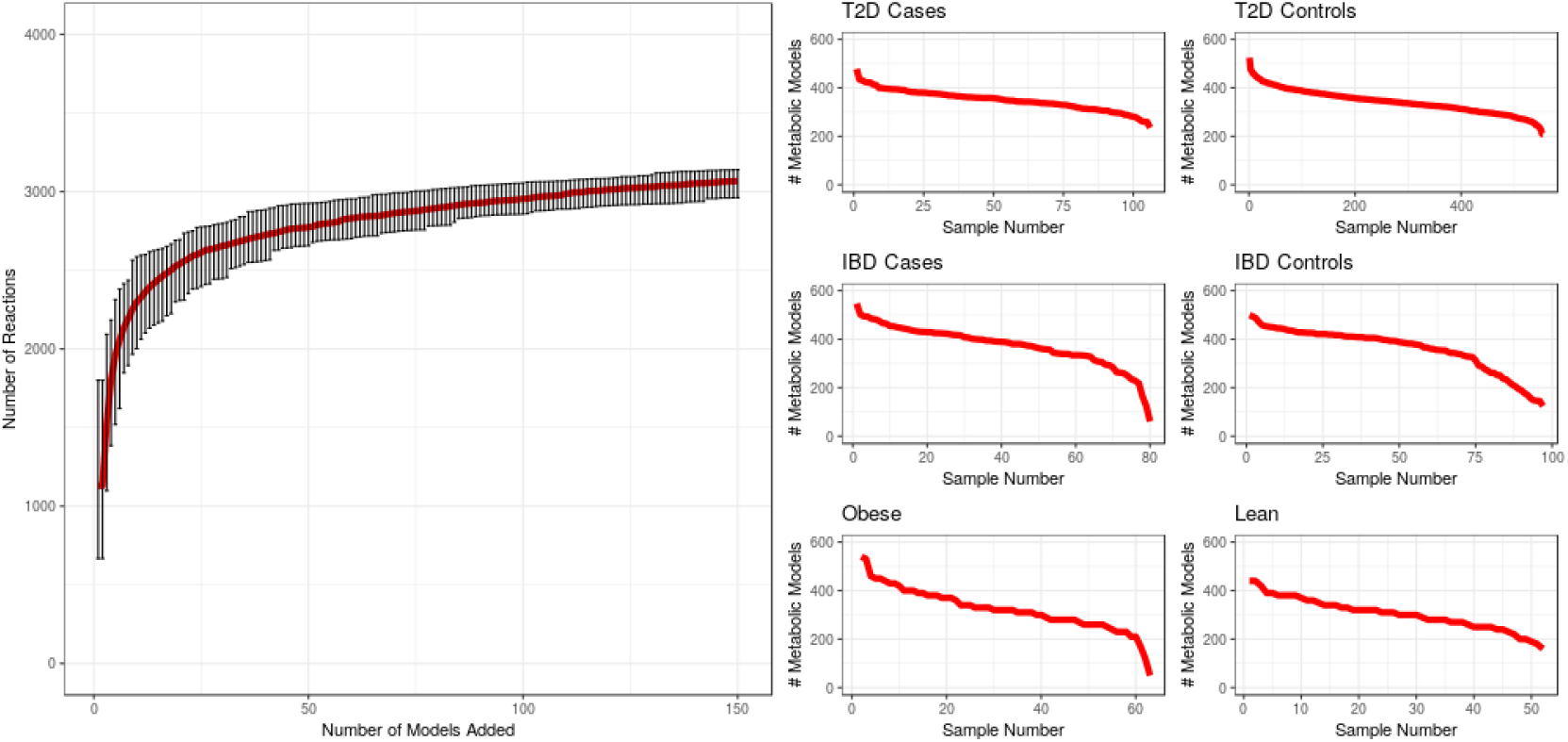
Left: For 1000 iterations, individual AGORA models were added to merged model in randomized order. Median, minimum, and maximum number of reactions in the merged model, after adding up to 150 individual models, are shown. Right: Number of individual metabolic models that were found to be present in each sample, from the three microbiome studies that were examined in this paper.

Finally, in order to map metagenomic reads for each species to determine their abundance, we followed two different procedures among our three disease-specific datasets. For both IBD and diabetes datasets, we downloaded appropriate tables of species abundances, based on 16s reads, from their supplementary data. For our obesity dataset, only metagenomic samples were obtained, and such a table was not available, so we used the following pipeline to extract species abundances. First, we first filtered for 16S reads in each metagenomic sample of the obesity dataset using sortmerna [27]. We ran this program with the following parameters: using the SILVA 16s database [28], and we requiring a 90% match threshold of reads to a 16s dataset. We then used the QIIME2 [29] command pick_closed_reference.py to map the filtered 16S reads, output by sortmerna, against the Greengenes database taxonomy [30]. This results in species abundances based on 16s reads from the obesity dataset.

With the 16S abundances of each species in each study, we then used PICRUST [31] to infer abundances of each species-specific metagenome, with the default parameters as given in the paper. These metagenomes were used in FALCON to simulate the metabolism of single gut microbiome species. For paired-model simulations, we used the inferred metagenomes of both modeled species. Finally, for merged model simulations, we added together the abundances of all inferred metagenomes. That is, for a reaction i in the merged model, we determined all individual models that contained reaction i, along with reaction i’s inferred metagenomic abundance in each individual model. We added together the inferred abundances of reaction i, and used that as the abundance of reaction i in the merged model. We used this method, as opposed to simply mapping metagenomic reads to the merged model, because all species in the gut microbiome would contribute to metagenomic reads, including some not included in the AGORA models. However, the merged model was initially created from individual AGORA models, and we only wish to include the metagenomic abundances of those individual AGORA models.

### 3.1 Calculation of Taxonomic Distances Between Individual-Species Models

We downloaded version 13.5 of the Greengenes taxonomy [32]. We matched the names of individual AGORA species models to leaf nodes of the taxonomy, and then calculated the taxonomic distance between every pair of such leaf nodes. We used these pairwise taxonomic distances to examine the correlation of metabolic cooperation/competition with taxonomic distance (see Pairwise Models section of Results).

## 4 Results

### 4.1 Merged and Individual-Species Models

We performed simulations on metagenomic data from several studies that examined three different metabolic diseases: obesity [23], T2D [24], and IBD [25]. Using the pipeline described in Methods, we extracted species-specific metagenomes with PICRUST. We then mapped expression from each of these three datasets onto two different types of metabolic models, first the set of all individual species models present in a sample, and then the merged model of all individual species models. This results in a total of six different cases, and we performed FALCON simulations on each of them. To determine which subsystems have differential flux between normal control and patient samples, we first analyzed individual reactions. We first ranked all reactions with differential flux, based on a Wilcoxon rank-sum test for differential flux between control and diseased groups, and determined all those reactions with a Wilcoxon p-value < .05 for differential flux. For each comparison between metabolic diseases and controls, there were hundreds of such nominally significant reactions, as we did not correct for multiple testing. It should be noted, though, that the actual flux difference of flux difference between the two groups was quite small for all reactions. For example, Figure 3 shows the distribution of fluxes between diabetics and normal controls for a very highly differential reaction in flux, phosphoglycerate kinase, showing a small but significant decrease of .00226 flux units (Wilcoxon p-value .021) in diabetics. Finally, we calculated the enrichment of each subsystem for these differential flux reactions, based on a hypergeometric test.

**Figure 3:**
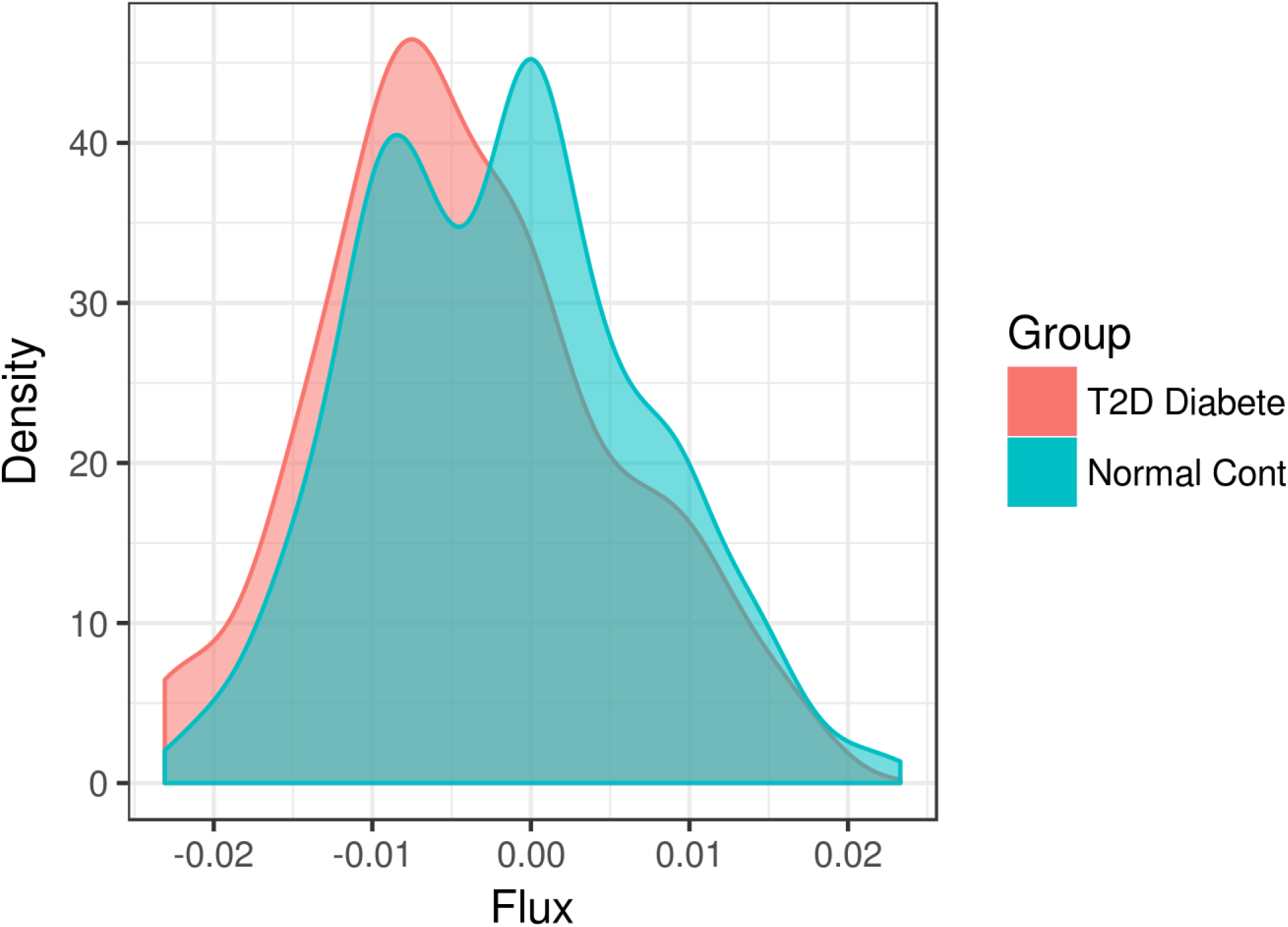
Distribution of fluxes for phosphoglycerate kinase, simulated using the merged model, among 106 T2 diabetic samples and 554 normal controls.

The results are summarized in Figure 4 below for the case of the merged model simulations. Several interesting findings emerge when we compare across the three diseases. The pentose phosphate pathway, lipopolysaccharide biosynthesis, and lysine, branched amino acid, and histidine metabolism are all down-regulated in expression across at least one dataset. These pathways are all related to biosynthesis in some way, and thus their downregulation in patients suggests that the gut microbiota grows at a slower rate. It is also interesting to note that branched chain amino acid synthesis is downregulated in T2D patients versus normal controls, as this supports evidence that shows higher levels of such acids in diabetic patient bloodstream. Our analysis suggests that the gut microbiome may be a contributor to this difference. It is known that gut microbiome metabolism can be extensively intertwined with that of the human host. Metabolites like butyrate can be exported from the microbiome to the host, where it serves as an energy source for colonocytes [33], and conversely nutients such as mucin can be provided by the host to the microbiome [34]. On the other hand, the expression of starch and sucrose metabolism and fatty acid oxidation is upregulated in some patients, suggesting increased use of these metabolites for energy.

**Figure 4:**
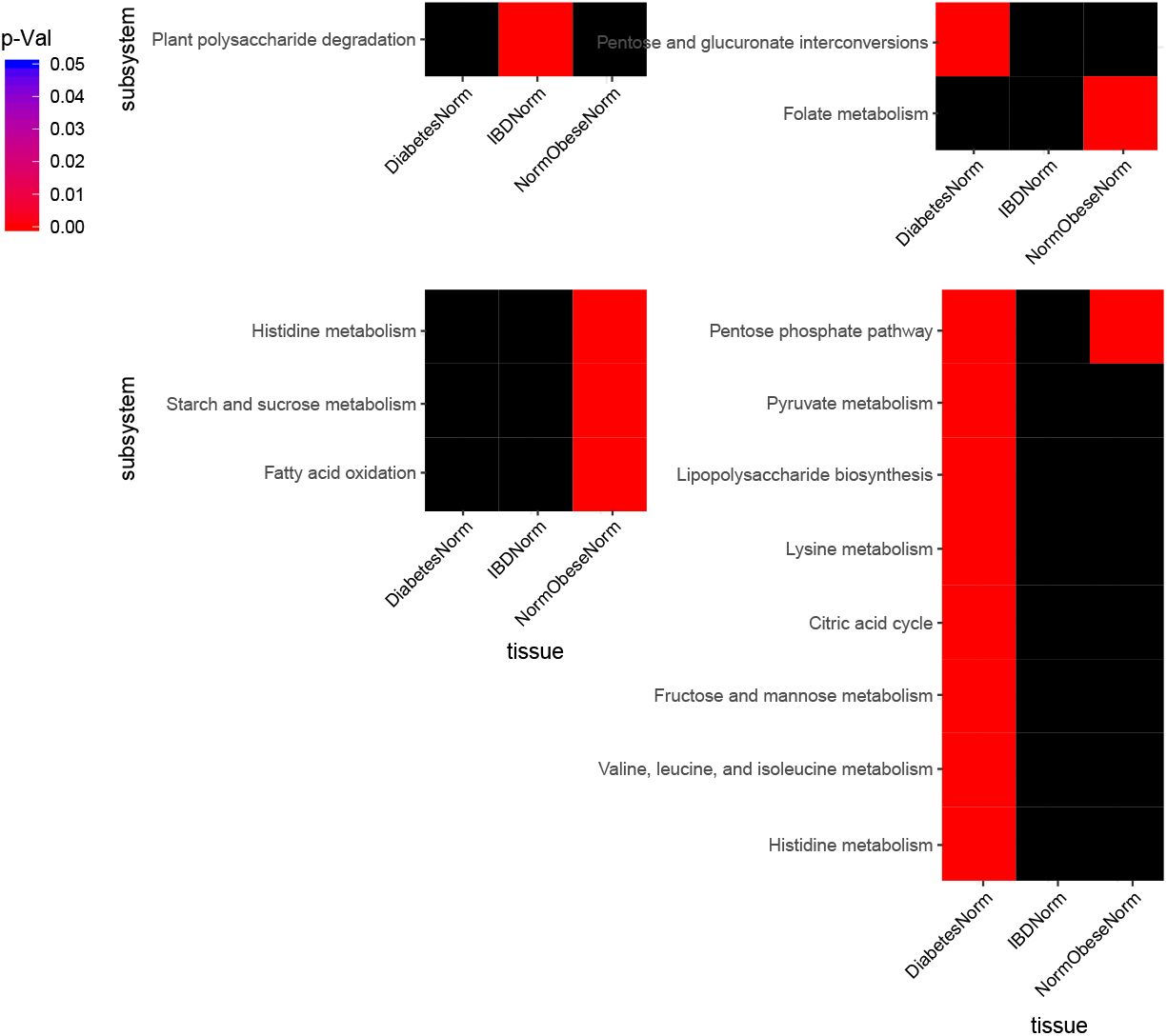
Top-left: Positive differential flux subsystems. Top-right: Negative differential flux subsystems. Bottom-left: Positive differential expression subsystems. Bottom-right: Negative differential expression subsystems.

Finally, significant differential flux pathways were fewer than differential expression pathways. Plant polysaccharide degradation was upregulated in IBD patients, while folate metabolism and glucuronate interconversion were down-regulated in obese and T2D patients respectively. Polysaccharide degradation is surprising to see as upregulated in patients, because it serves as a precursor to synthesis of SCFAs such as butyrate, which are associated with a healthy microbiome.

We were also interested in possible flux differences between the single-species AGORA models, vs. the merged model. We expect that flux predictions from the two types of models would be very different, as the merged model is able to capture all possible species interactions, compared to individual species models which cannot capture any. We therefore compared the predictions for differential flux between the merged and single-species models, using each of our three datasets. In general, there was very poor correlation of individual fluxes between the two simulation cases. The merged model is expected to be more accurate in this case because it can capture more species interactions.

The poor correlation between individual species and merged models can be seen in Figure 5. In these plots, we considered respectively reactions that showed either a positive or negative merged-model flux difference in normal control samples vs. patients. We also determined the corresponding reactions in the single-species simulations, and the sum of either positive or negative individual species fluxes for each reaction. Then, we correlated merged-model positive differential flux with individual species positive differential flux, and similarly for negative differential flux. As can be seen, in Figure 5, there does not seem to be a significant correlation between the two quantities in any dataset. This indicates that differential flux in the merged model does not have a significant association with individual species differential flux.

**Figure 5:**
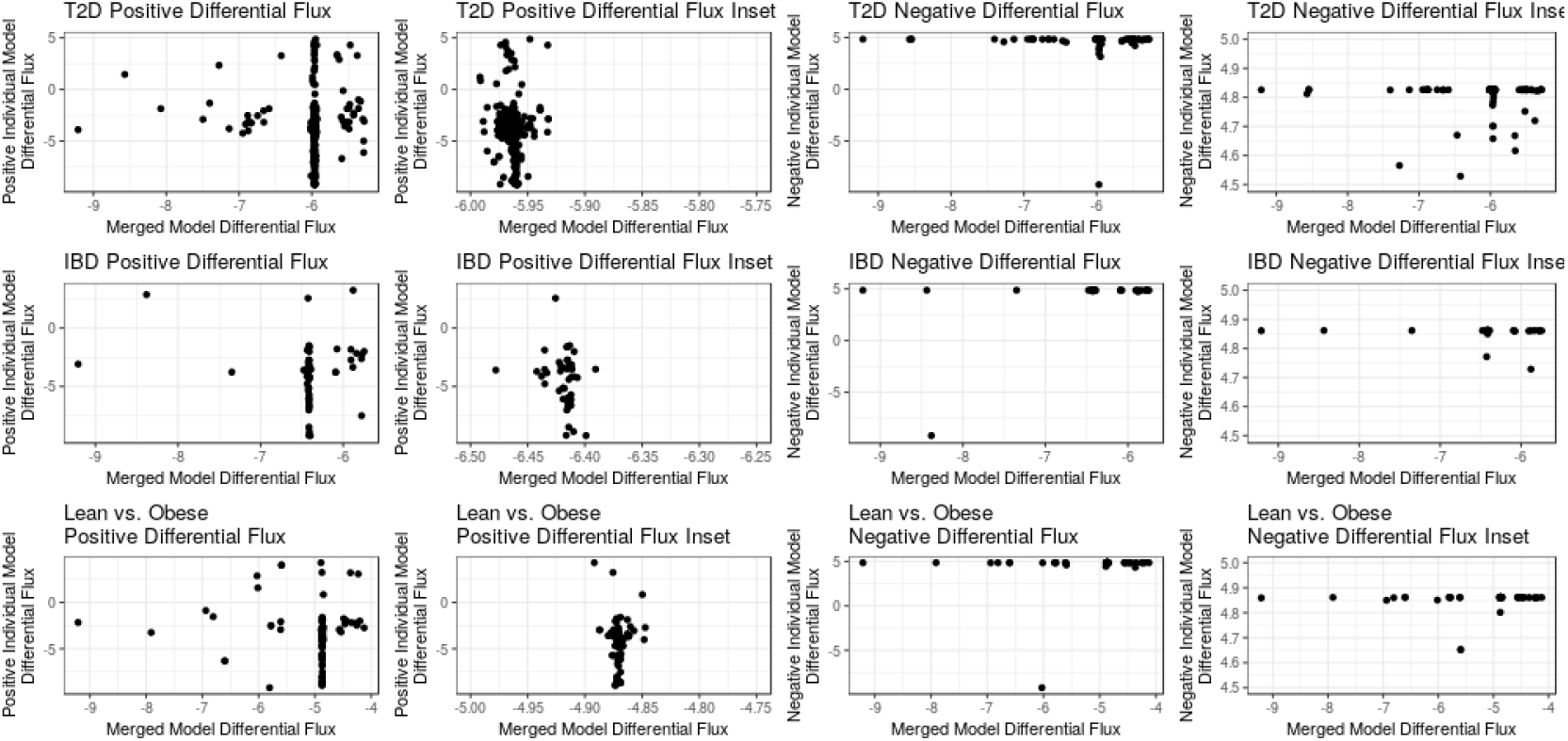
Average merged model differential flux, vs. average individual species differential flux, across samples from the three studies that were examined. Inset panels are focused on areas where data points were clustered together, in order to a get a clearer view.

An even more fine-grained analysis is shown in Figure 6, which consider only reactions that showed significant differential flux in both the merged model and single-species cases. In this case, the x-axis of the heatmap indicates the number of reactions in each dataset with either positive or negative merged model differential flux (Wilcoxon p-value < .05). On the y-axis is shown the individual species log differential flux, normalized to range between −1 to +1, and colored green for positive differential flux, red for negative, and black for no significant difference. If there were any association between merged-model and individual species differential flux, we would expect to see a trend in which positive merged-model differential flux is associated with more positive individual species differential flux, and vice versa. However, the plots in Figure 6 do not show any such trend.

**Figure 6:**
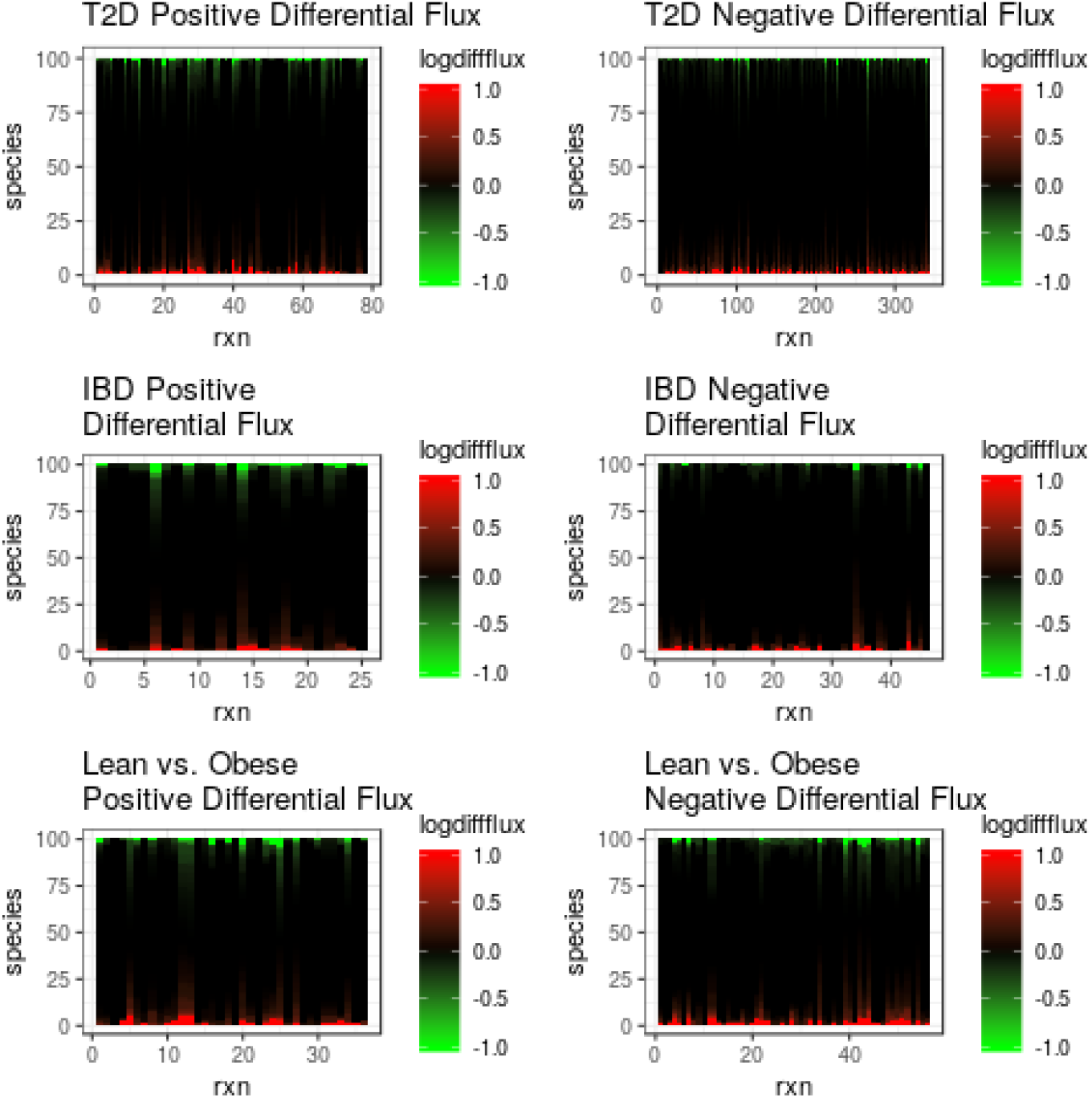
For reactions with significant merged model differential flux in each dataset, we plotted either positive (green) or negative (red) log differential flux in individual species.

Furthermore, there was also poor correlation of the differential flux when comparing any of the three metabolic disease datasets against each other, as shown in Tables 1 and 2. These results suggest that the overall metabolic changes between the three metabolic diseases we study are very divergent.

**Table 1:** Correlations between differential flux predictions among three metabolic diseases vs. controls, for single-species models

| Comparison Type | Spearman's R | Spearman's R-squared | Spearman's p-val |
| --- | --- | --- | --- |
| (IBD vs. normal)<br>vs. (T2D vs. normal) | -.06 | .0035 | .0012 |
| (Obese vs. normal)<br>vs. (T2D vs. normal) | .135 | .018 | 7.64E-12 |
| (Obese vs. normal)<br>vs. (IBD vs. normal) | -0.019 | .00038 | .33 |

**Table 2:** Correlations between differential flux predictions among three metabolic diseases vs. controls, for merged models

| Comparison Type | Spearman's R | Spearman's R-squared | Spearman's p-val |
| --- | --- | --- | --- |
| (IBD vs. normal)<br>vs. (T2D vs. normal) | -.017 | .00029 | .39 |
| (Obese vs. normal)<br>vs. (T2D vs. normal) | -.026 | .00068 | .141 |
| (Obese vs. normal)<br>vs. (IBD vs. normal) | -.035 | .00123 | .07 |

Finally, Figure 7 shows the average abundance levels and metabolic flux levels of all reactions in the three datasets plotted against each other. It shows there is very poor correlation between the level of enzyme abundance in the metagenomic data, versus the level of predicted flux. Therefore, the nature of gut microbiome metabolic changes cannot be predicted from expression alone, but instead must be based on simulated flux measurements, using methods like FALCON.

**Figure 7:**
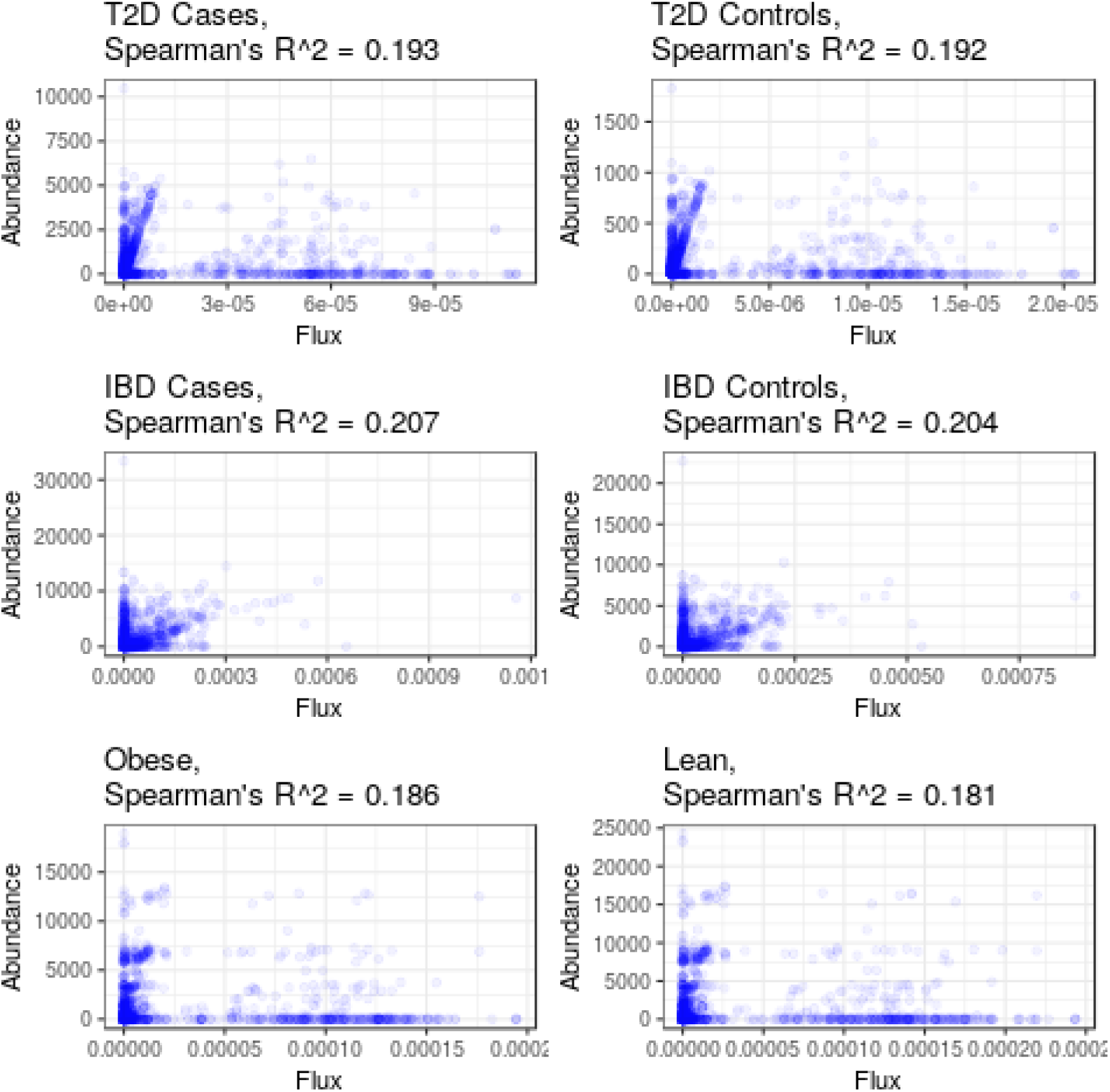
Average abundance level of reactions versus average predicted flux, across all three datasets.

### 4.2 Pairwise Models

Our main interest in pairwise FALCON simulations is to measure the degree of metabolic cooperation or competition in the gut microbiome. We therefore adapted a previously published metric by Sung et al. [35] Their formula was designed to compare the number of metabolites that are both uptaken as nutrients by a pair of species, versus those metabolites that are shared by two species via crossfeeding. We modified this formula, to take into account the magnitude of fluxes involving such metabolites. Our formula for the influence of species i upon species j, through metabolic cooperation/competition, is thus

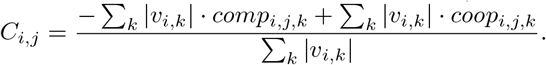

In this equation, the index *k* runs over all metabolites that can be either uptaken or excreted by species *i. v_i,k_* represents the magnitude of the uptake or excretion flux for metabolite *k* in species *i*. Finally, the coefficient *comp_i,j,k_* is 1 if metabolite *k* is either uptaken by both i and *j*, or released by both of them, and 0 otherwise. Similarly, *coop_i,j,k_* is 1 if metabolite *k* is released by species *i* and uptaken by *j*, or vice versa, and 0 otherwise. The matrix *C* can therefore be for pairwise flux simulations involving any metagenomic sample, and represents the set of net metabolic cooperation/competition for all pairs of species.

Previous work on metabolic cooperation in the gut microbiome [35] has suggested there is a negative correlation, between the degree of cooperation in a species pair, versus the taxonomic distance between a species pair, as measured by the Greengenes taxonomy [32]. Species pairs that are taxonomically distant are expected to differ more in their metabolic network structure and reactions. This should reduce the chance that two species both compete for the same nutrients, and also increases the probability that they crossfeed for a certain metabolite. However, surprisingly, we find that there is no correlation between the degree of metabolic cooperation and taxonomic distance among species pairs, as shown in Figure 8. One possible interpretation of this result is that species at all taxonomic distance actually face equal evolutionary pressure to metabolically cooperate and/or compete with each other. In that case, the magnitude of metabolic flux may be adjusted, so that overall cooperation is equal at all taxonomic distances.

**Figure 8:**
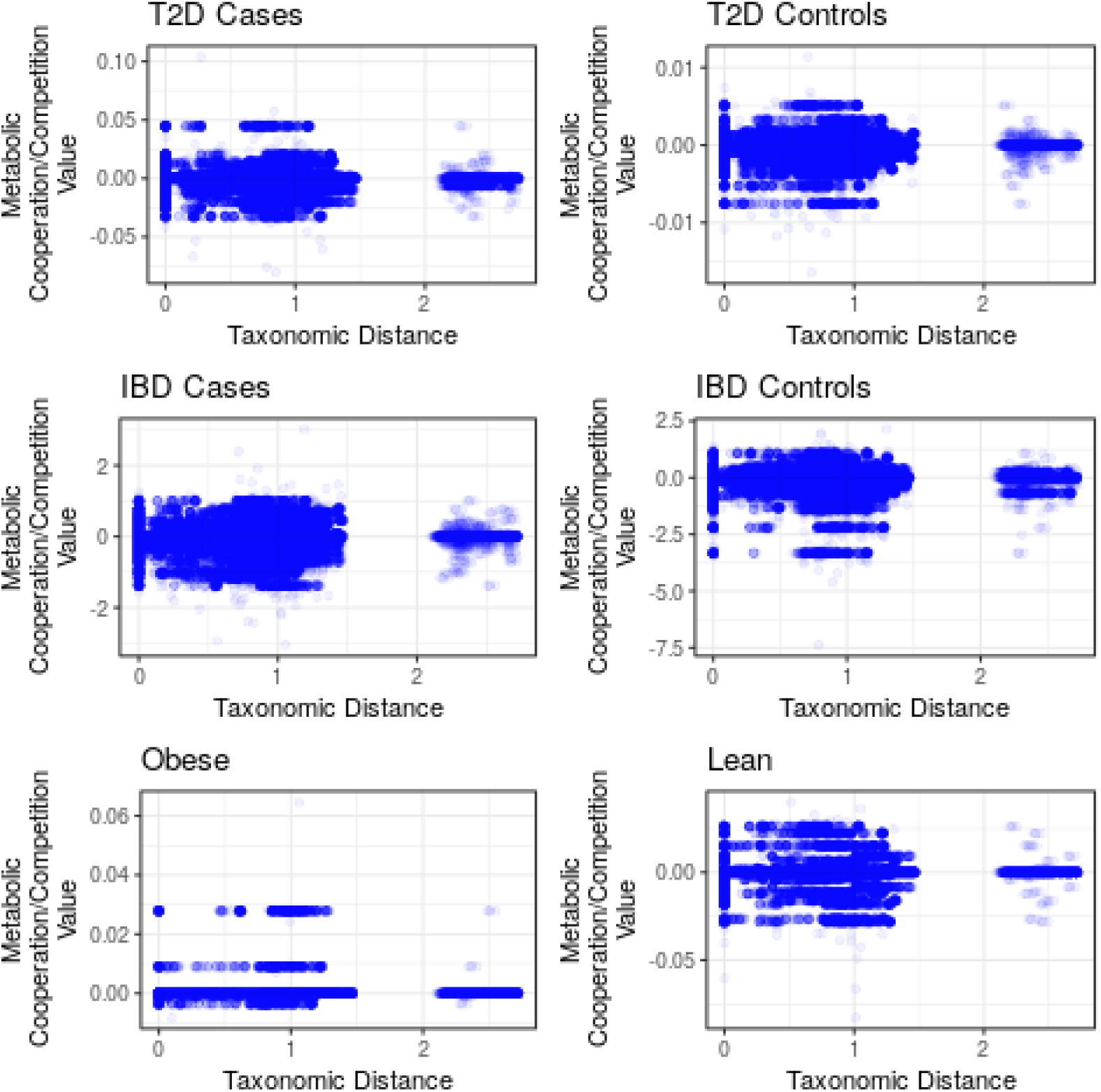
Level of metabolic cooperation vs. taxonomic distance among species pairs, in all three datasets. Note that the smaller cluster to the right of the figure represents taxonomic distances measured between archaeal and bacterial species, and the cluster to the left represents distances between two bacterial or two archaeal species. Archaeal-bacterial taxonomic distances are expected to be considerably greater than between members of the same domain.

From this matrix of cooperation/competition values, we then investigated the compositional stability of the gut microbiome. Compositional stability is one of the most important properties of the microbiome, as a host must have a stable microbiome in order to count on benefits from it. Our work here is inspired by a recent study from Coyte et al. [36] Their mathematical framework begins with a matrix *A*, where *A_i,j_* represents the influence of species *j* on growth of species *i*. The values of A are not known experimentally, but can be sampled from a distribution of interaction types. Positive values indicating a beneficial effect of species *j*’s abundance on species *i*, and negative values a detrimental effect. The abundance and dynamics of species *i*, symbolized by *X_i_*, is then given by:

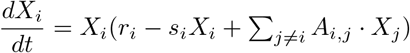

where *r_i_* represents the intrinsic growth rate of species *i*, and *s_i_* the carrying capacity of species *i*. The eigenvalues of this matrix are then calculated and used to determine stability, as explained later in this paper.

The matrix *C* that we defined earlier has two major differences from *A*. Both stem from the fact that in all FALCON simulations in this study, we were unable to simulate flux through a biomass reaction, and therefore could not calculate growth rate directly. Firstly, we therefore could not determine the *r_i_* and *s_i_* coefficients found in the above formula, and must set them to 0 in our calculations. Secondly, whereas the coefficients of *A* represent the direct effects of species *j* on growth of species *i*, our matrix *C* represents this effect indirectly. As described before, our *C* contains the weighted average of metabolites under either competition or cooperation between *i* and *j*. Nevertheless, as metabolite uptake and excretion is known to be a major constraint on microbial growth rates [19], we believe our metric does capture at least some features of cooperation and competition. Therefore, we calculated cooperation/competition matrices for the datasets we described previously, and calculated eigenvalues to determine their stability.

Eigenvectors and eigenvalues are a concept from linear algebra that can be used to determine the sample’s compositional stability. For any n x n square matrix, a set of n eigenvectors may be calculated, and each eigenvector is associated with one eigenvalue. The set of eigenvectors for the matrix spans all possible directions of change in the sample’s composition, so that any perturbation in composition can be represented as a linear combination of eigenvectors. Furthermore, the direction in which the composition moves, after a perturbation, is determined by the eigenvalues. Negative eigenvalues indicate that the system will move back to its original starting point after a perturbation in the direction of an eigenvector. Positive eigenvalues indicate that the system will continue moving away from the starting point, in the direction of the perturbation. Therefore, only if all eigenvalues of the cooperation/competition matrix are negative, will the composition of the system be stable.

The top ten eigenvalues, by absolute value, for both normal controls and patients in all of our three datasets, are plotted in the figures below (Figure 9). For all of our three datasets, there were some positive eigenvalues in both normal control and patient samples, indicating that they were all unstable. However, this result is not necessarily unexpected, as all negative eigenvalues imply the **entire** microbiome is stable, so that no single species are ever lost or gained. This is actually contradicted by data such as [37], which found that a few species are lost or gained over a period of 1 year in one subject. Furthermore, we infer that the time-scale of such changes may correlate with the magnitude of the largest positive eigenvalue for each sample. From linear algebra, it is known that for a larger positive eigenvalue, a system will move more quickly away from it’s original starting point, in the direction given by the corresponding eigenvector. Thus, for the T2D and IBD comparisons, the magnitude of the largest positive eigenvalue was greater in control samples compared to patients, indicating that control samples actually change more quickly than patients in these two cases. However, the opposite is true in lean samples vs obese, with lean samples having greater magnitude of the largest positive eigenvalue.

**Figure 9:**
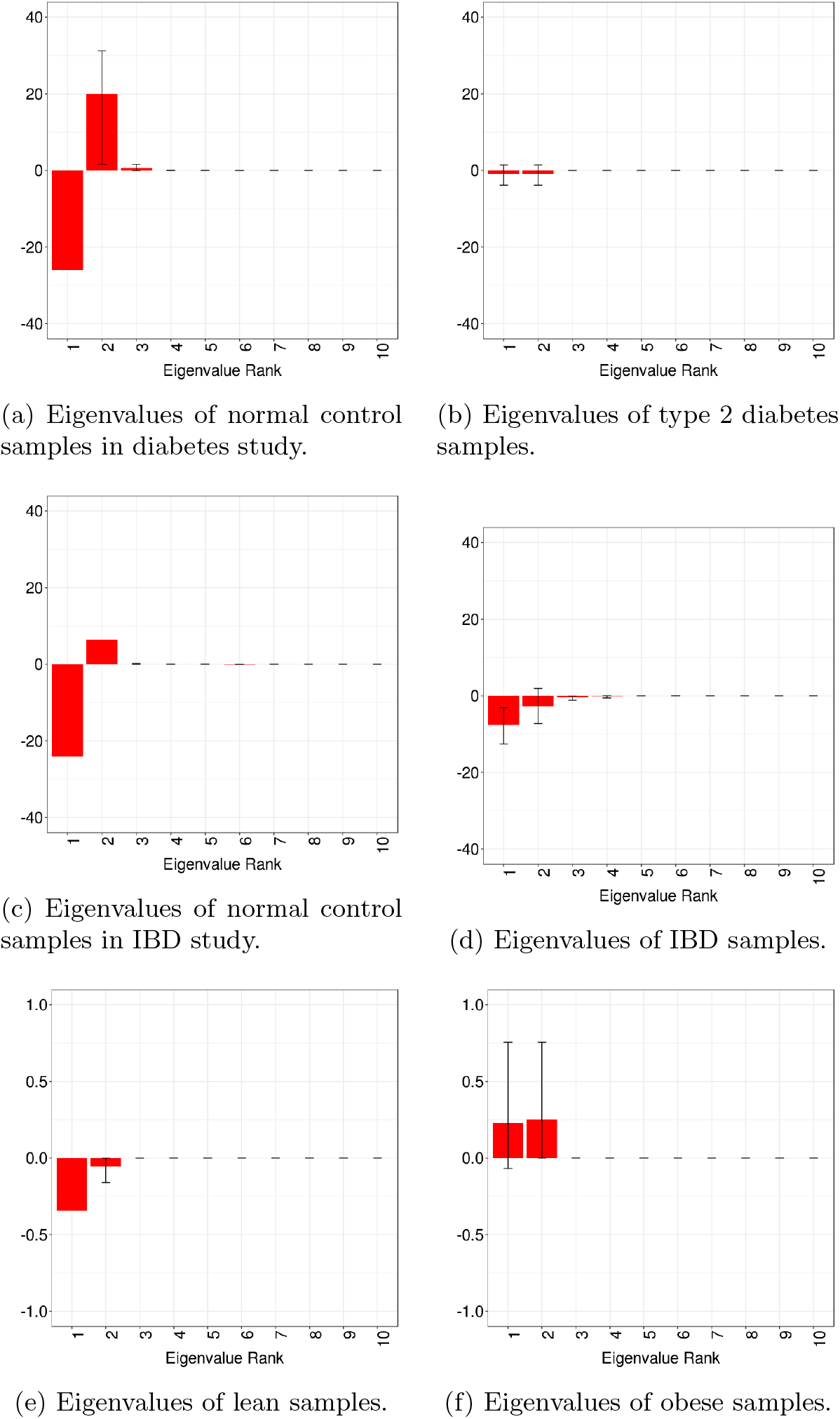
Eigenvalues of pairwise models for diabetic, IBD, and obese samples, versus respective controls.

These results were unexpected for us, as previous observations suggest approximately equal stabilities of healthy and diseased microbiomes [38]. Our interpretation is that control samples in the T2D and IBD datasets do not occupy a single stable species composition, but rather change constantly within a finite space of compositions, which are consistent with having a healthy microbiome. T2D and IBD patients would also move within a finite space, corresponding to a disease state, but at a slower rate than healthy samples. Similarly, lean samples would be predicted to move within their compositional space at a slower rate than obese samples. Since 2 out of 3 metabolic diseases in this study thus are associated with slower gut microbiome compositional changes compared to controls, further studies may show whether or not this is a general trend among all microbiome-associated diseases.

## 5 Discussion

The problem of inferring metabolic flux distributions is a key problem in computational biology, and is essential to unraveling the mechanisms of many complex metabolic diseases. So far, most work in this area has focused on single bacterial species, but the gut microbiome is made up of hundreds of species, which together play a major role in the metabolism of the human host. To predict flux in this challenging environment, we modeled the gut microbiome with three separate models, either individual-based models, pairwise models, or a merged model, as described in Methods. We then applied a gene-expression based method, called FALCON, to infer flux distributions for metagenomic samples from either normal controls, or patients with T2D, IBD, or obesity.

Using this approach, we first identified metabolic pathways that had significant differential flux between controls and patients. Somewhat surprisingly, we observed that the overall correlation between differential metabolic flux in each of the three metabolic diseases is surprisingly low. These three metabolic diseases have high co-occurrence with each other, and are often considered to share similar metabolic causes and outcomes in the human host, and thus possibly the gut microbiome as well. However, our results suggest that, when looking at metabolism as a whole, which includes many pathways outside of central carbon metabolism, there may be little similarity between them. This may be relevant in the future, as new approaches that target novel metabolic pathways for treatment of one disease, may not apply to the others.

Nevertheless, at the level of pathways, we found that all three diseases have a few pathways with a very significant differential flux, and that a few of these are shared among these diseases. Among these is branched amino acid synthesis, which has previously been shown to be involved in human host metabolic changes. Our results suggest that the microbiome as well may contribute to disease-specific metabolic changes, and thus may be a promising target for therapies, as several available probiotics already set out to do.

Another important question in the gut microbiome is the extent of metabolic cooperation or competition among species. Interactions among species help to determine the overall composition of the gut microbiome, as well as its stability in the face of perturbations. Here, our results were unexpected, as by using the eigenvalues of the matrix of interactions, we found that there are some positive eigenvalues in nearly all samples, indicating that all samples are unstable. Although this result is different from previous work [36], we believe there is some justification for it, based on the observation that real microbiomes do slowly gain and lose species over time [37]. Furthermore, our results show that the time-scale at which microbiome compositions change is different for metabolic disease vs. control, with T2D and IBD patients changing more quickly than controls, while obese samples change more slowly than lean.

In conclusion, our study is an attempt to model the metabolism of the gut microbiome, using gene-expression based modeling, and how it may contribute to common metabolic diseases in humans. Our results give insight into how the gut microbiome may affect common metabolic symptoms and mechanisms in these diseases. We expect that future studies may further improve the accuracy of metabolic inference in the gut microbiome.

